# Detection of cross-frequency coupling between brain areas: an extension of phase-linearity measurement

**DOI:** 10.1101/2020.11.25.398628

**Authors:** Pierpaolo Sorrentino, Michele Ambrosanio, Rosaria Rucco, Joana Cabral, Leonardo L. Gollo, Michael Breakspear, Fabio Baselice

## Abstract

The current paper proposes a method to estimate phase to phase cross-frequency coupling between brain areas, applied to broadband signals, without any a priori hypothesis about the frequency of the synchronized components. N:m synchronization is the only form of cross-frequency synchronization that allows the exchange of information at the time resolution of the faster signal, hence likely to play a fundamental role in large-scale coordination of brain activity. The proposed method, named cross-frequency phase linearity measurement (CF-PLM), builds and expands upon the phase linearity measurement, an iso-frequency connectivity metrics previously published by our group. The main idea lies in using the shape of the interferometric spectrum of the two analyzed signals in order to estimate the strength of cross-frequency coupling. Here, we demonstrate that the CF-PLM successfully retrieves the (different) frequencies of the original broad-band signals involved in the connectivity process. Furthermore, if the broadband signal has some frequency components that are synchronized in iso-frequency and some others that are synchronized in cross-frequency, our methodology can successfully disentangle them and describe the behaviour of each frequency component separately. We first provide a theoretical explanation of the metrics. Then, we test the proposed metric on simulated data from coupled oscillators synchronized in iso- and cross-frequency (using both Rössler and Kuramoto oscillator models), and subsequently apply it on real data from brain activity, using source-reconstructed Magnetoencephalography (MEG) data. In the synthetic data, our results show reliable estimates even in the presence of noise and limited sample sizes. In the real signals, components synchronized in cross-frequency are retrieved, together with their oscillation frequencies. All in all, our method is useful to estimate n:m synchronization, based solely on the phase of the signals (independently of the amplitude), and no a-priori hypothesis is available about the expected frequencies. Our method can be exploited to more accurately describe patterns of cross-frequency synchronization and determine the central frequencies involved in the coupling.

## 1. Introduction

Brain areas need to constantly transfer information among themselves to put in place complex behavioural responses to the environment [1]. Functional connectivity is defined as the presence of statistical dependencies between the time-series representing the activity of brain regions [2, 3]. A variety of mechanisms through which this communication occurs are summarized in [4], involving only the phase [5] or also amplitude [6]. Each of these phenomena would underlie a specific neuro-physiological mechanism (for a review, see [7]). In the literature, a wide number of metrics have been proposed to detect each of these kinds of communication [8, 9]. Furthermore, communication between brain areas can occur either in iso-frequency or in cross-frequency. Cross-frequency coupling (CFC) is the interaction occurring between neuronal populations operating at different frequencies. It has been postulated that this form of synchronization could represent a suitable option to allow large-scale synchronizations across distant areas in the brain [10, 11], yielding the integration of distributed information [12]. Moreover, definite (both frequency and spatial) patterns of CFC have been shown to be the neuro-physiological substrate underlying the recruitment of areas needed for the execution of tasks such as specific kinds of learning [13, 14, 15], segregation of interfering inputs [16], perception [17, 18], encoding of reward [19] or sensory processing [20]. In human brain activity, two main forms of cross-frequency coupling have been described so far. Firstly, the phase of slow oscillations can modulate the amplitude of faster activity [21, 22]. Furthermore, phase-phase synchronization has also been described, whereby the phases of “n” cycles of a signal are locked to “m” phase cycles of another signal [5]. This kind of cross-frequency communication, classically defined as n:m synchronization, has been observed previously in human brain data [23] and is the only mechanism capable of supporting CFC at high temporal resolution [24].

Several metrics have been developed to capture the presence of cross-frequency communication. For instance, phase-amplitude coupling [9] can successfully detect the presence of nested-synchronization, while metrics such as bicoherence [25] can detect cross-frequency, phase-phase coupling. However, bicoherence is not a pure phase-based metrics as its value depends also on the amplitude, preventing an unambiguous interpretation of the involved neuronal mechanisms [22]. The biphase-locking value, while purely based on the phase, also provides an estimate of the phase-amplitude coupling [26]. Metrics such as the phaselocking factor [22] detect pure phase to phase locking, but require an accurate a priori hypothesis about the frequencies involved in cross-frequency synchronization. The procedure proposed by Cohen [27], on the other hand, while not requiring any a priori hypothesis, focuses on phase-amplitude coupling.

Each approach has its own advantages and drawbacks and, when one is dealing with specific task-related data, given that a specific a priori hypothesis is available about the frequencies across which synchronization might be occurring, the application of these metrics is effective [12]. However, when dealing with resting-state data, the situation changes because the signals contain several frequency bandwidths interacting with each other (possibly with more than one of the mentioned mechanisms) at once.

Restricting the analysis to phase-to-phase coupling, we have to consider that the bandwidth of the involved signals is so broad and complex to potentially allow the simultaneous occurrence of iso- and cross-frequency synchronizations at once [12, 6, 11, 28]. Disentangling these complex signals has proven to be elusive when one does not know a priori if, when and where cross-frequency is occurring within the brain. The aim of this paper is to develop a reliable estimation of phase-phase cross-frequency communication between the broadband signals of two brain regions, without a priori hypothesis on the frequencies at which such a synchronization might occur. To do this, we build and expand upon the phase linearity measurement (PLM), an iso-frequency phase-based connectivity metrics recently developed by our group [29].

One issue is related to the amount of potential combinations of frequencies and areas that one should test in order to look for CFC throughout the brain and throughout the frequency spectrum. Indeed, an attempt to identify the frequency at which cross-frequency synchronization is present from the data by selecting a number of combinations of possible frequencies has been done [30],using the level of synchronization across trials in order to statistically estimate where cross-frequency synchronization was present.

To this regard, a new method has been recently proposed, that does not require any a priori hypothesis and can estimate cross-frequency synchronization [31]. Such an approach estimates from the data the “candidate frequencies” where the CFC might be occurring. However, when performing this procedure, a maximization of the correlation between the signals is performed, hence reintroducing a form of dependency from the amplitude. The issue of the communication between different frequencies has also been addressed using a multiplex network approach [32]. The idea is that each layer of the multiplex network represents, at a specific frequency, the iso-frequency correlations between brain areas. However, the cross-layers links are not estimated from the data. With the methodology proposed in this paper, we aim at providing a data-informed estimate of which brain areas and frequencies are involved in cross-frequency phase-to-phase coupling. The novelty of this work lies in the fact that no a priori information is required about the frequencies and the areas involved in the CFC. On the contrary, our technique allows to start from wide signal spectrum and to detect if cross-frequency is occurring and, if so, to identify which frequency components are involved per each signal. Firstly, we provide a theoretical description of the metric. Secondly, the metric is tested in a number of synthetic analytical models. We first used Rössler oscillators, which capture the non-linearities of the brain. Secondly, in order to simulate the simultaneous presence of iso and cross frequency synchronization, we implemented several Kuramoto oscillators, and introduced a lag between the generated signals. This procedure is known to produce the appearance of synchronization at a lower frequency bandwidth as compared to the original signals [33]. Hence, we tested the ability of the newly proposed methodology, namely cross-frequency phase linearity measurement (CF-PLM), to detect and disentangle both kinds of synchronism. Furthermore, we mixed the previously produced signals linearly, in order to obtain a case in which both iso-frequency and cross-frequency coupling are simultaneously present, and we tested if the newly proposed approach can disentangle such a situation. Finally, we tested the metrics on source-reconstructed MEG data (acquired by the MEG laboratory in Naples), and identified brain areas where cross-frequency is present that are spatially consistent across the tested subjects.

## 2. Methods

### 2.1. Definition of the interferometric signal

Let us define *x*(*t*) and *y*(*t*) as the time series related to two brain areas. By applying the Hilbert transform, their analytical expression is obtained:

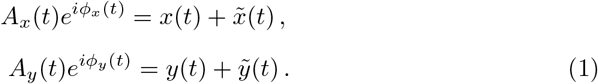

where variables *A* and *ϕ* represent the amplitude and the phase, respectively. According to this mathematical description, signals generated by brain areas can be modeled as complex phasors with time-varying amplitude and phase. According to [29], their phase-to-phase connectivity can be measured via a three steps procedure. Firstly, one extracts the normalized interferometric component of the two signals *z*(*t*) by computing:

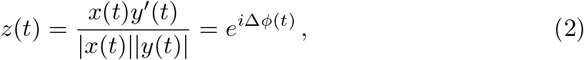

where the symbol ′ indicates the complex conjugate. Note that the complex interterometric function *z*(*t*) has an amplitude equal to 1 (thus it is independent of the amplitudes of the signals *A*_*x*_(*t*) and *A*_*y*_(*t*)), and a phase term Δ*ϕ*(*t*) = *ϕ_x_*(*t*) *− ϕ_y_*(*t*) *∈* [*−π, π*[, which is the time-varying phase difference between the phases of *x*(*t*) and *y*(*t*). It has been shown that one can exploit the behavior of the term Δ*ϕ*(*t*) in order to measure phase connectivity between signals. Hence, the frequency analysis of the function *z*(*t*) is carried out. Three different conditions could occur, as reported in Figure 1. In case of no synchrony between the sources, the interferometric phase values appear to be irregularly spread in the [*−π, π*[range (blue line in Figure 1). In case of phase coupling, the term Δ*ϕ*(*t*) will be characterized by a linear trend. That is, if the two sources have a similar oscillation frequency, the phase of the interferometric signal will be constant or slowly varying in time (red line in Figure 1) while in case of two sources oscillating at different frequencies, a slope will appear (yellow line in Figure 1). Once the complex signal *z*(*t*) has been computed, the second step for measuring the coupling consists in computing its power spectrum by means of the Fourier transform:

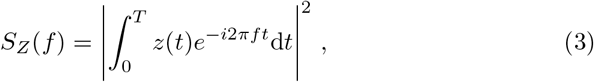

where [0, *T*] is the observation period. In order to have a more reliable evaluation of the PSD function, we implemented the periodogram estimator with a rectangular window and confidence interval of 0.95 for the computation of *S_Z_*(*f*) [34]. The shape of the power spectrum is strongly influenced by the strength of the coupling occurring between the two sources and by their central frequency, and hence it can be exploited to estimate them [29].

**Figure 1:**
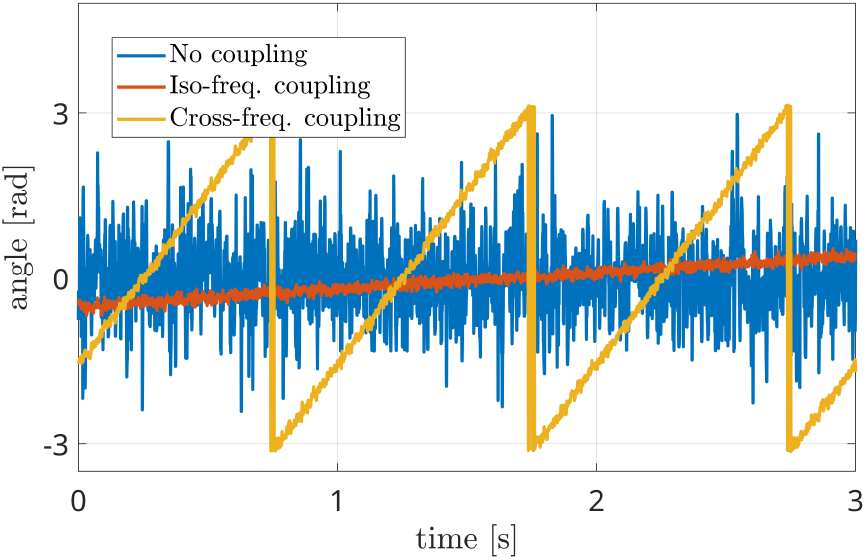
Interferometric phase signals in three different conditions: independent sources (blue line), coupled iso-frequency sources (red line) and coupled sources at different frequencies (yellow line).

### 2.2. Phase Linearity Measurement

In Figure 2, the power spectra occurring in the different scenarios are represented. The blue line does not show any peak, in accordance with the absence of coupling between the sources. This means that the power spectrum of *z*(*t*) is almost flat (blu line of Figure 2) if its phase term Δ*ϕ*(*t*) irregularly spreads in the [*−π, π*] range (blue line of Figure 1).

**Figure 2:**
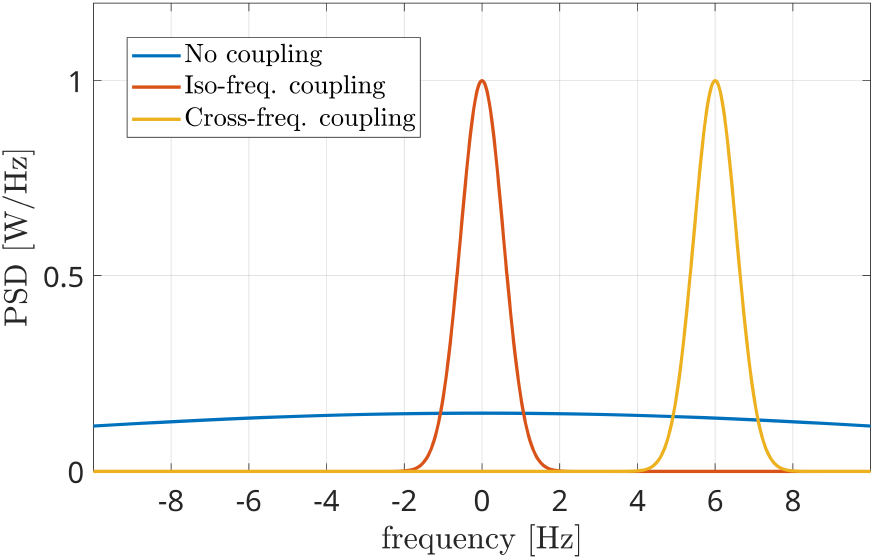
Power spectral densities of the interferometric components(i.e. the power spectrum of the phases of the interferometric signal) in three different conditions: independent sources (blue line), coupled iso-frequency sources (red line), coupled sources at different frequencies (yellow line). The presence of a power peak denotes the coupling between sources, while its position indicates the difference in their resonant frequencies.

The red line shows an evident power peak around *f* = 0, which is due to a linear behavior of the interferometric phase Δ*ϕ*(*t*), i.e.:

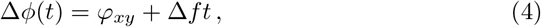

where the term Δ*f* is related to the different resonance frequencies of the two sources. In the case of iso-frequency coupling (IFC), such a term is relatively small, resulting in a peak centered around *f* = 0. In this case, the last step for measuring connectivity strength consists in computing the percentage of power within a narrow band [*−B, B*] around *f* = 0:

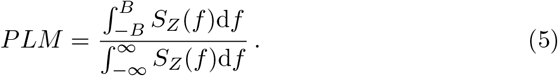

In [29] it has been shown that a *B* value of 1 Hz is a well balanced trade-off between the discrimination capability and the estimation noise of the algorithm. The PLM approach has shown a good performance in measuring the iso-frequency coupling, i.e. in distinguishing between the case of the blue line and the red line in Figure 2. Nevertheless, it has to be modified in order to make it effective in analyzing the last case, the cross-frequency coupling.

### 2.3. Cross-frequency PLM

In the CFC condition, a non-minimal frequency difference Δ*f* occurs between the coupled components of the sources, and such difference produces a shift in the interferometric spectrum, as shown by the yellow curve of Figure 2. In this case, the coupling is evident due to the presence of the peak, which is now centered at *f* = Δ*f* (6 Hz in the reported case) instead of *f* = 0. This difference makes the PLM (Eq. (5)) unable to capture the coupling, as the power is no longer concentrated in the [*−B, B*] band. One should notice that the knowledge of the frequency difference Δ*f* would solve the problem, as the integration could be shifted accordingly into the [Δ*f − B,* Δ*f* + *B*] frequency range. However, this kind of *a priori* knowledge is not available at all times, as it is often the case in resting-state as well as in many task-related settings. This situation can be handled by looking for maxima (i.e. power peaks) in the interferometric power spectrum *S_Z_*(*f*). Once a local maximum is identified (besides the one centered in 0), its power and position can be easily measured. This is what the proposed methodology implements. In other words, once the PSD function of Eq. (3) is computed, the global maximum is identified. By retrieving its position, the difference between the two sources central frequencies Δ*f* is identified. Subsequently, the coupling strength is measured by adapting the upper integral of Eq. (5), i.e.:

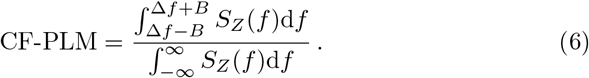

However, the provided information concerns the frequency difference between the two sources, the central frequencies of the coupled components still have to be determined. Hence, the last step is to identify the oscillation frequencies of the two components involved in the CFC. To this aim, a band-stop Gaussian-shaped frequency filter has been adopted. The stop band is centered at *f_H_* and is 2*B* large. The central frequency *f_H_* is moved in order to scan the whole frequency range of the acquired signals, e.g. the [0.5, 48] Hz range, as reported in Figure 3 (top). Let us focus on the signal of the first source (i.e. *x*(*t*) of Eq. (1)). Once the filter is overlapped to frequency components involved in the coupling and removes them, the peak of the interferometric PSD *S_Z_*(*f*) disappears, as shown in Figure 3 (center). The filter position *f_H_* will reveal the frequency *f_x_*of *x*(*t*) involved in the coupling. The same process is repeated for the second source *y*(*t*) for the identification of *f_y_*, according to Figure 3 (bottom). After the frequency scans, the two central frequencies of the components involved in the CFC *f_x_* and *f_y_* are identified, while the amount of coupling is related to the peak energy and is measured via Eq. (6).

**Figure 3:**
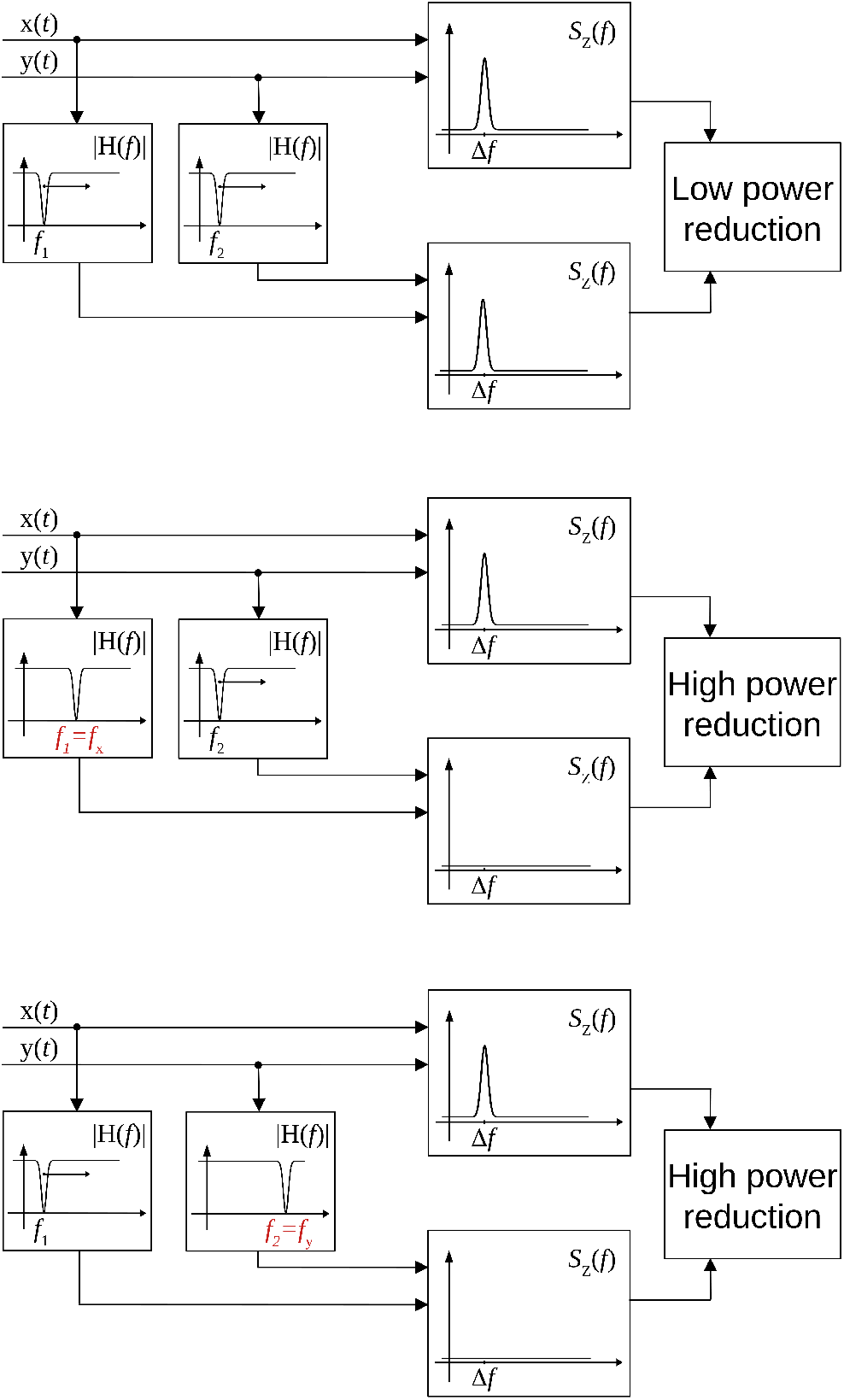
Scheme of the procedure for the identification of frequencies involved in the coupling. When the frequency stop filters are not overlapped to the frequencies involved in the coupling, the peak in the interferometric PSD is present (top). When one filter overlaps with the frequency of the first (center) or the second source (bottom), the peak disappears and there is a reduction in the power.

## 3. Results

The proposed methodology has been tested on both synthetic and real datasets. In case of simulated data, two approaches have been adopted for generating the cross-frequency coupled signals, exploiting Rössler attractors and Kuramoto oscillators, respectively. In more detail, the sensibility of the CF-PLM metrics to coupling strength has been analyzed by means of Rössler attractors signals to which a frequency shift has been applied. Furthermore, a modified version of the Kuramoto oscillators implementing signals with different central frequencies has been considered, in order to test the ability of the CF-PLM to identify the two frequencies involved in the coupling. As a third analysis, real data acquisitions have been considered for the final validation of the approach.

### 3.1. Rössler attractors

Two time series have been generated according to the model [35]:

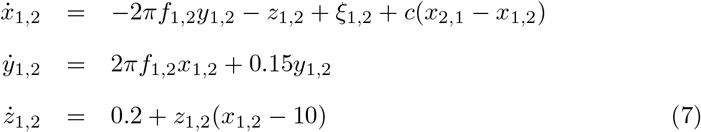

with coupling strength *c* varying between 0 and 0.04. A resonance frequency *f*_1_ equal to 10 Hz has been chosen, while the duration and the sampling interval have been set equal to 625 Hz and 420 s, respectively. The two coupled time series *x*(*t*) and *y*(*t*) have been generated with a central frequency *f*_1_. Subsequently, the cross-frequency has been simulated by applying a frequency shift to the second attractor *y*(*t*) via the modulation property of the Fourier transform:

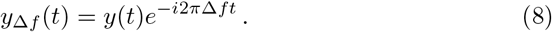

For this analysis, we considered Δ*f* = 7 Hz. The CF-PLM has been computed between *x*(*t*) and *y*_Δ*f*_ (*t*). Several analyses have been conducted aiming at evaluating the sensitivity of the proposed metrics with respect to the coupling strength of the attractors, the SNR and the frequency shift. In Figure 4, the values measured by the CF-PLM as a function of attractor’s coupling strength are reported in case of different SNR levels (using white, additive noise). In more detail, a Monte Carlo simulation with 50 iterations has been implemented and the mean values are reported. As expected, the CF-PLM value increases as a function of the coupling strenght, for each of the considered noise levels. Moreover, we tested the CF-PLM in case of several frequency shifts, obtaining the same curves reported in Figure 4.

**Figure 4:**
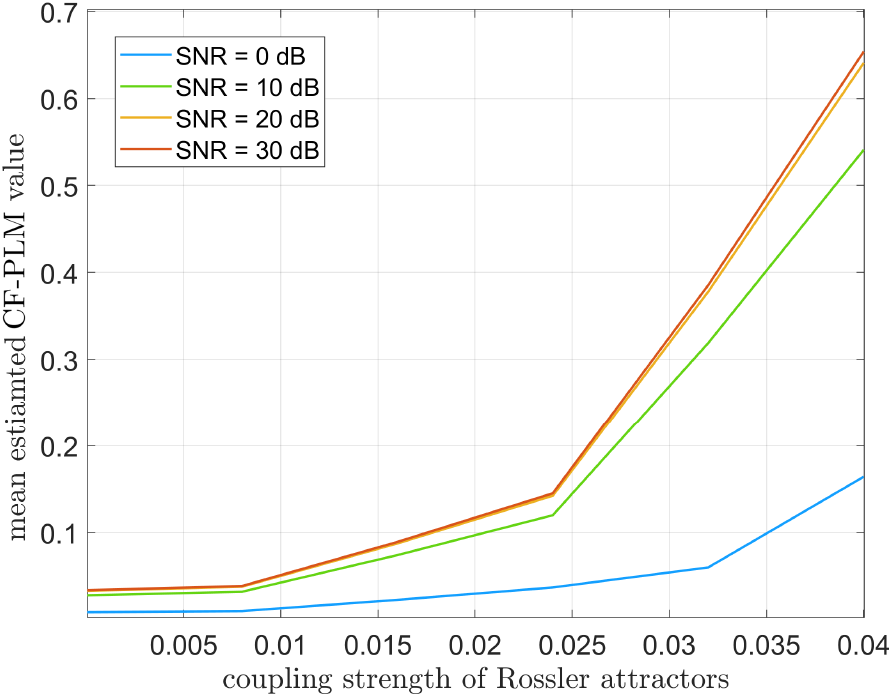
Mean values of CF-PLM measured in case of two Rössler oscillators varying their coupling strength from 0 (no coupling) to 0.04 (high coupling). Results are reported in case of different SNR values between 0 dB and 30 dB.

### 3.2. Kuramoto oscillators

Three mutually coupled Kuramoto oscillators, namely *s*_1_(*t*), *s*_2_(*t*) and *s*_3_(*t*), have been generated according to the following model [36]:

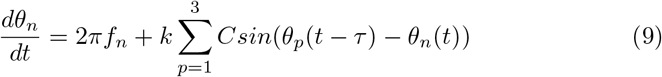

with *n* = 1, 2, 3, *τ* = 0.6 s and *C* = 1. The central frequencies have been set equal to 10 Hz, 10 Hz and 17 Hz, respectively. The first oscillator (*s*_1_(*t*)*, f* = 10 Hz) has been compared to the second one (*s*_2_(*t*)*, f* = 10 Hz), to the third one (*s*_3_(*t*)*, f* = 17 Hz) and to the sum of the last two (*s*_2_(*t*) + *s*_3_(*t*)*, f* = 10 Hz and 17 Hz). The three PSDs of the interferometric signals *S_Z_*(*f*) are reported in Figure 5. It is evident that iso-frequency synchronization is correctly measured in the case of two 10 Hz oscillators (the power peak centered at 0 Hz of Figure 5a), as well as the cross-frequency synchronization occurring between *s*_1_(*t*) and *s*_3_(*t*) (the power peak centered around −7 Hz of Figure 5b). Importantly, the case of multiple components simultaneously synchronized in iso and cross-frequency is correctly handled, with the two power peaks positioned at 0 Hz and −7 Hz in Figure 5c.

**Figure 5:**
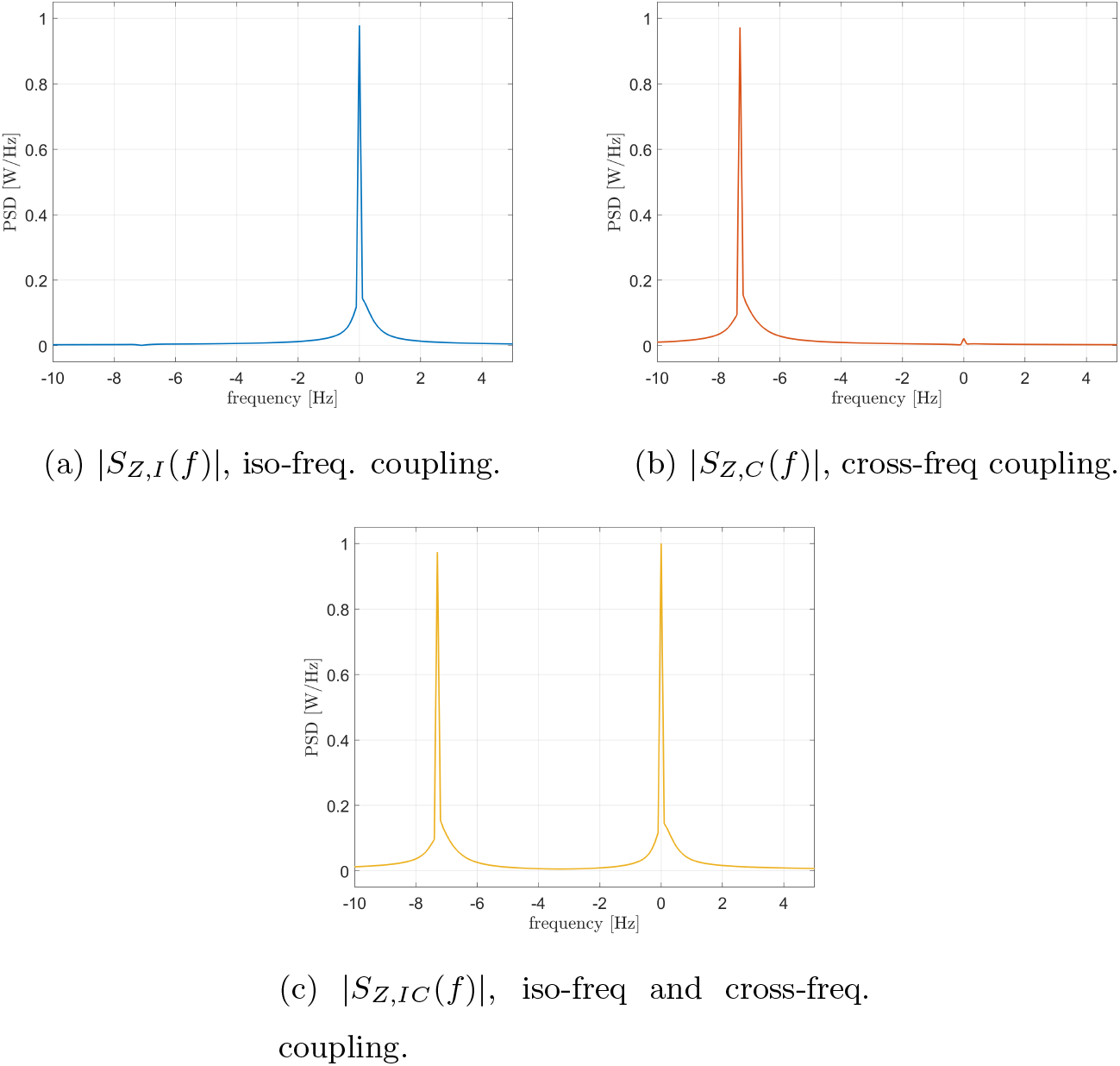
Power spectra of the interferometric signals in case of coupled Kuramoto oscillators: iso-frequency (a), cross-frequency (b), iso and cross-frequency (c).

Since the Kuramoto oscillators are coupled with a time delay between them, the frequency shift depends not only on the natural frequencies of each oscillator but also on the coupling strength between them [37, 38]. In Figure 6 we show that the position of the interferometric peak is shifted as a function of the coupling strength, thus validating the existence of a cross-frequency interaction between the oscillators.In other words, this shows that the presence of synchrony, at the frequency that is predicted theoretically, is captured by the metric (as opposed to merely be capturing n:m phase relationships, whose frequencies are not expected to be dependent from the coupling strength).

**Figure 6:**
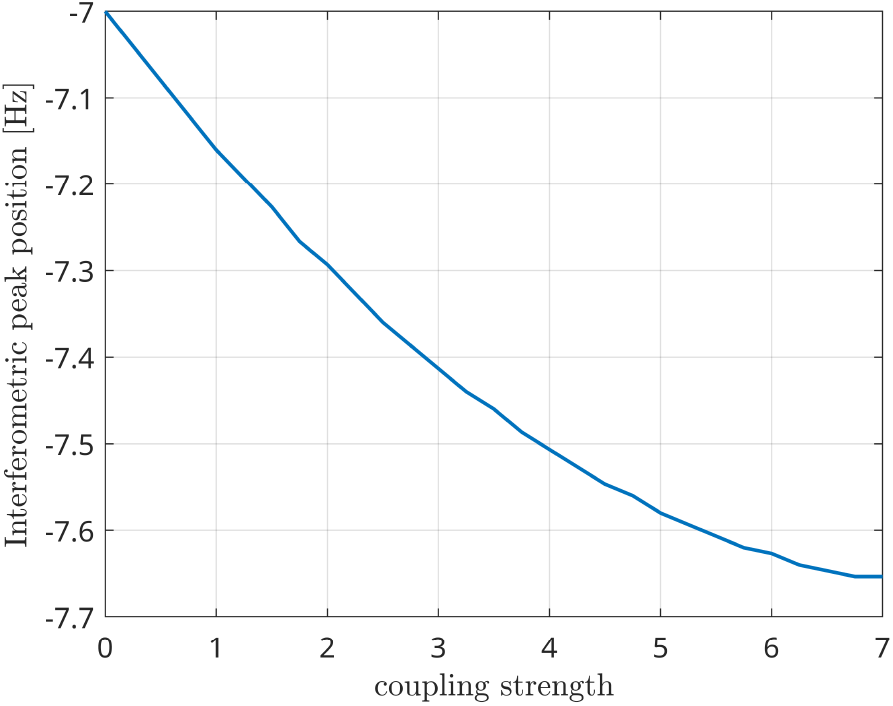
Position of the interferometric peak while varying the global coupling strength of two Kuramoto oscillators.

Let us now analyze how the frequencies involved in the connectivity process are identified. According to the processing scheme previously described and reported in Figure 3, two stop-band filters are implemented in the frequency domain. The peak power reduction is computed when moving the central frequencies of these filters within the [0, 20] Hz range. Results are reported in Figure 7 for all the considered cases. When the filter removes from the first signal the frequencies involved in the coupling, a power reduction is measured in the PSD peak. As a consequence, a horizontal dark line will appear in the images of Figure 7. Analogously, a vertical line will appear when the corresponding frequency of the second source is removed. The result is a cross-shaped image, with the center identifying the two frequencies involved in the coupling. By looking at Figure 7, it is evident that the maximum power reduction appears at (10, 10) Hz in the case of *s*_1_(*t*)*, s*_2_(*t*) coupling, at (10, 17) Hz in the case of *s*_1_(*t*)*, s*_3_(*t*) coupling, while in the simultaneous iso and cross-frequency coupling two couples are correctly identified at (10, 10) Hz and (10, 17) Hz, respectively. All these results are in accordance with what we expected, as the procedure correctly estimates both the connectivity strength and the oscillator frequencies involved in the coupling from the interferometric spectrum.

**Figure 7:**
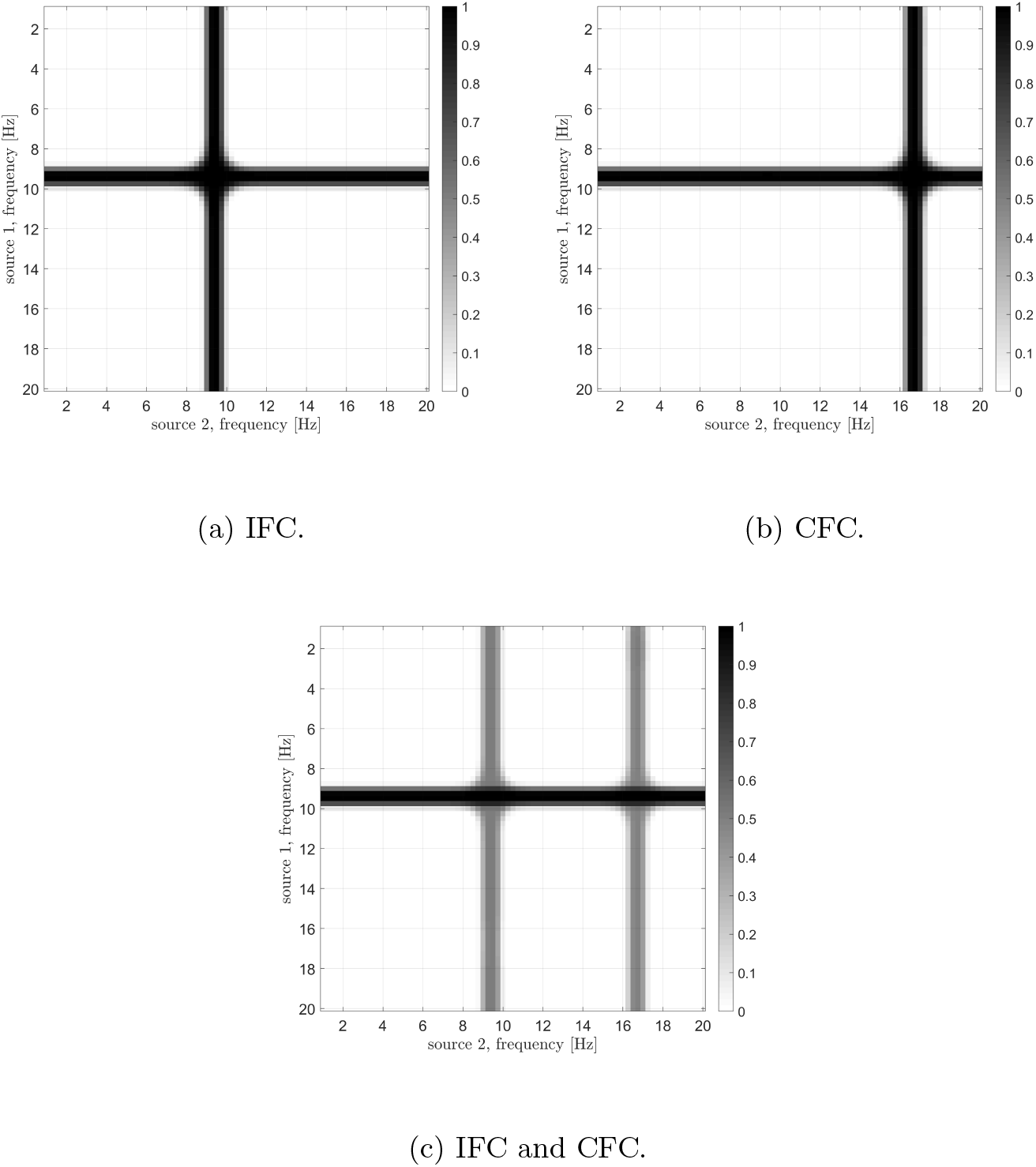
Results of the analysis for the identification of frequencies involved in the coupling in case of different Kuramoto oscillators: iso-frequency (a), cross-frequency (b), and simultaneous iso- and cross- frequency (c). The center of each cross identifies the frequencies of the two oscillators

In order to have a benchmark, the dual-frequency coherence (DFC) [39], which is a normalized version of the second order bispectrum [40], has been implemented. Given the two acquired signals *x*(*t*) and *y*(*t*) and their Fourier transform *X*(*ω*) and *Y* (*ω*), the DFC is defined as:

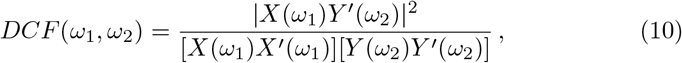

We computed the DFC for *ω*_1_ and *ω*_2_ in the [1, 20] Hz range in case of the CFC Kuramoto oscillators, obtaining the result reported in Figure 8. Compared to Figure 7b, it is evident that the metrics is less effective in determining the frequencies involved in the coupling as two maxima are present at a distance of about 1 Hz.

**Figure 8:**
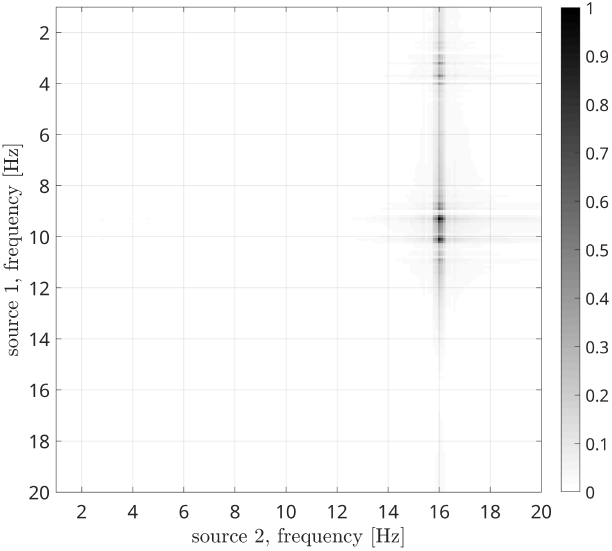
Dual-frequency coherence results in case of CFC Kuramoto oscillators.

### 3.3. Real data

#### 3.3.1. Acquisition and preprocessing

The acquisitions used for the analysis are from healthy subjects acquired at the MEG facility in Naples. The detailed procedure used for the processing of the acquisitions has been described in [41].

In brief, subjects were seated in a 163-magnetometers MEG system placed in a magnetically shielded room (AtB Biomag UG - Ulm - Germany). The brain activity was recorded twice for 3.5 minutes, with a small break to minimize the chances of drowsiness. After the anti-aliasing filter, the data were sampled at 1024 Hz, and filtered between 0.5 and 48 Hz with a 4^th^ order Butterworth IIR band-pass filter [42]. During the acquisitions, the electrocardiogram (ECG) and the electrooculogram (EOG) were also recorded [43]. Principal component analysis (PCA) was used to reduce the environmental noise [44, 42]. Subsequently, noisy channels were removed manually through visual inspection by trained experts. For each subject, supervised independent component analysis (ICA) [45] was performed to eliminate the ECG and, if present, the EOG components from the MEG signals. MEG data were then co-registered to the native MRI of the subjects. We used the volume conduction model proposed by Nolte [46] and the linearly constrained minimum variance (LCMV) beamformer [47] to reconstruct the time-series related to the centroids of 90 regions-of-interest (ROIs), derived from the automated anatomical labeling (AAL) atlas [48, 49, 50]. For each source, we projected the time series along the dipole direction explaining most variance by means of singular value decomposition (SVD), obtaining a scalar value per each source.

#### 3.3.2 Connectivity measurement

The power spectra of the interferometric signal for each couple among the 90 sources have been computed.

In the following, we selected two couples of regions, i.e. the couple with the highest CFC peak (among all couples of regions), and a region with an average CFC peak intensity. To show that a high CFC peak is unlikely to appear by chance, we have validated the analysis against surrogates. Hence, the intensity of the highest CFC peak, derived from 10000 random surrogates, obtained by shuffling the phase of the signal in the frequency domain, have been computed. In Figure 13 (a), we show that, in the case of the high observed cross-frequency peak, the peak intensity was above the 99th percentile of the surrogates. On the contrary, in case of the average CFC peak (Figure 13 (b)), its intensity was around the 50th percentile, as expected.

To further check the validity of our analysis in real subjects, we went on to estimate the anatomical consistency of the CFC peaks per each link across subjects. To do so, we proceeded as follows: for each of 4 subjects, for each source pair the intensity of the strongest CFC peak has been measured (|Δ*f| ≥* 2 Hz). Subsequently, we binarized the CFC peak matrix according to a threshold. To avoid a dependency of the result from the choice of the threshold, we have repeated the analysis across a range of thresholds, i.e. from the 1st to the 99th percentile. Then, we have summed across the binary matrices obtaining one matrix per subject which does not depend on any specific threshold (Fig. 13, a-d). Finally, the subject specific matrices were summed, showing that the presence of a CFC peak is topographically consistent across individuals (fig.13, e). In Figure 11 panel A, the source couples that are beyond a high number of thresholds, i.e. high percentile values, are reported in progressively intense red. The most evident points of this connectivity matrix are those related to the strongest CFC peaks. In panel B, the delta-frequencies corresponding to the significant edges are reported. Finally, in panel C, the regional sum of the peak intensity is reported.

Now we are going to focus on the couple of regions with the highest CFC peak. In Figure 9a, the PSD of the interferometric signal related to the right inferior parietal lobule and the orbital part of the right superior frontal gyrus are reported. Results are related to one epoch about 150 seconds long of a single subject. A power peak positioned at 9.5 Hz is clearly visible, which shows the presence of cross-frequency coupling. in the following step the sources frequencies involved in the coupling are determined. As explained earlier, the procedure consists in filtering the two signals and measuring the power reduction of the frequency peak. In Figure 9b, it is evident that the highest reduction is found in case of *f*_1_ *≈* 11 Hz and of *f*_2_ *≈* 1.5 Hz. The result obtained by the DCF, which is reported in Figure 9c, is not effective in identifying the involved frequencies, as the global maximum is hardly distinguishable.

**Figure 9:**
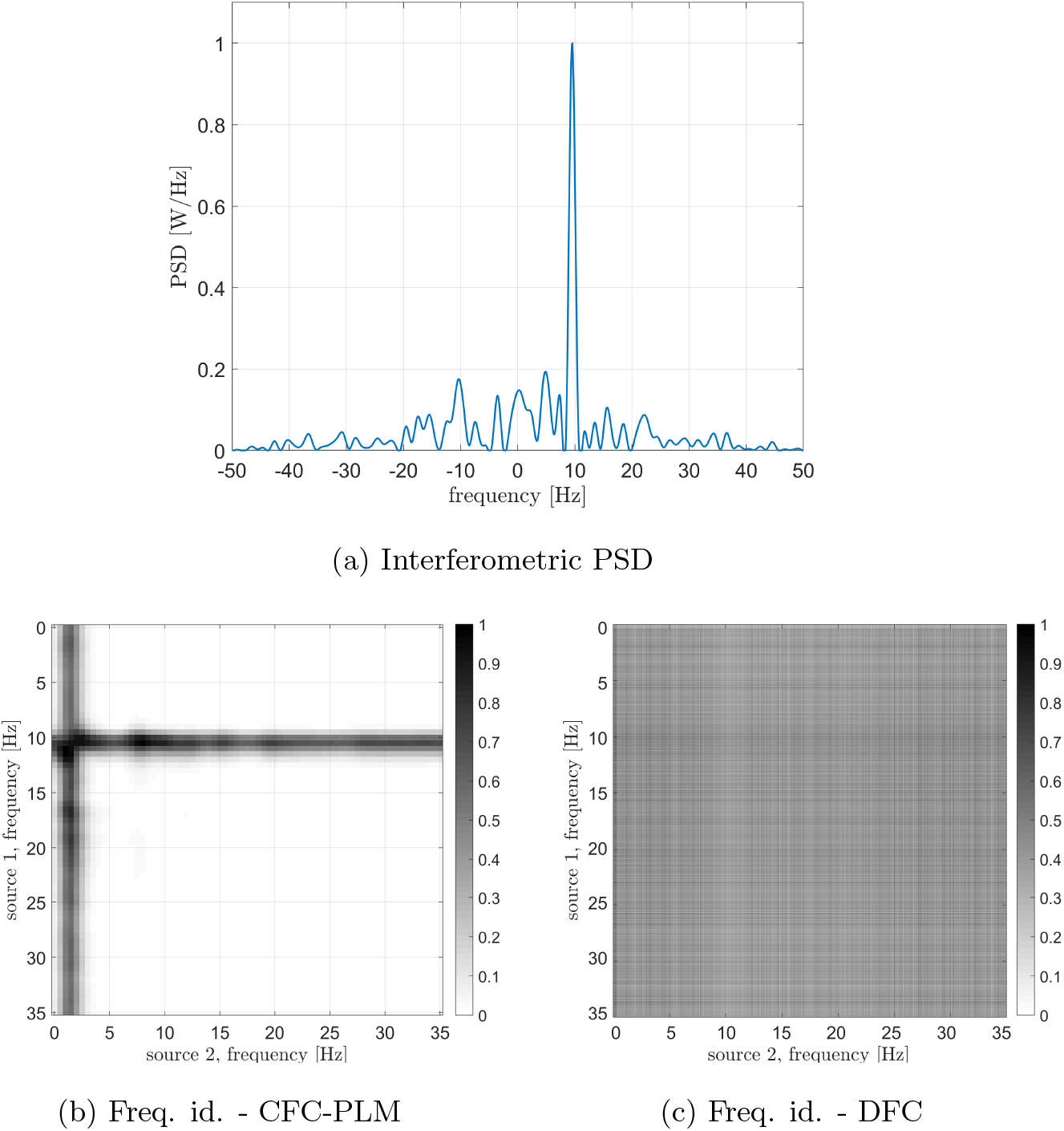
Analysis of signals from areas 57 (right inferior parietal lobule) and 42 (orbital part of the right superior frontal gyrus). (a) PSD of the interferometric signal. The peak related to the cross-frequency coupling, located at +9.5 Hz is clearly visible. (b) Frequencies identification via CFC-PLM. The frequencies involved are *f*_1_ = 11 (for source 57) and *f*_2_ = 1.5 (for source 42). (c) Frequencies identification via Dual-Frequency Coherence. Although a dark area in case of *f*_1_ = 10 Hz is visible, the two frequencies cannot be identified.

A second pair of brain regions has been considered, namely the left superior frontal gyrus and the left calcarine sulcus. The PSD of the interferometric signal is reported in Figure 10a. Two peaks are evident in this case, one centered in zero, related to the iso-frequency coupling, and another one at −8 Hz, which denotes a cross-frequency connectivity. By focusing on the latter, the identified involved frequencies are reported in Figure 10b, and are around 1 Hz and 9 Hz. Also in this case, results are more convincing than the DCF (Figure 10c), which cannot be exploited for the frequency identification.

**Figure 10:**
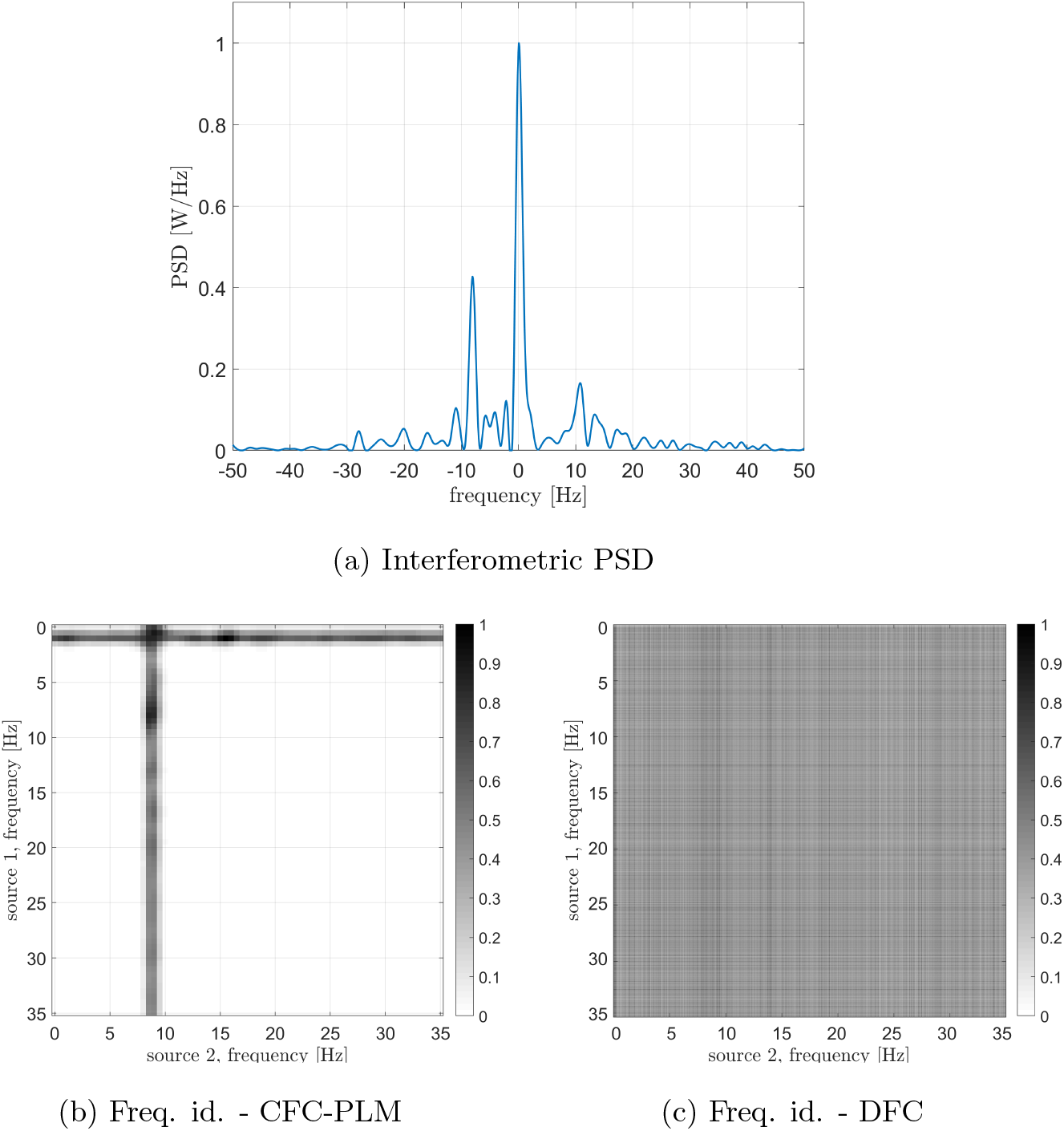
Analysis of signals from areas 7 (left superior frontal gyrus) and 25 (left calcarine sulcus). (a) PSD of the interferometric signal. Two main peaks are visible, located at 0 Hz (related to the iso-frequency coupling) and −8 Hz (related to the cross-frequency coupling). (b) Frequencies identification via CFC-PLM. The frequencies involved are *f*_1_ = 1 (for source 7) and *f*_2_ = 9 (for source 25). (c) Frequencies identification via Dual-Frequency Coherence. A global maximum is not visible, and thus the two involved frequencies cannot be identified.

## 4. Discussion

In this paper, we present a novel phase-based metrics capturing the occurrence of cross-frequency synchronization in the resting-state brain. The main advance of this work lies in the fact that the proposed procedure detects cross-frequency synchronization reliably, without a priori hypothesis about the frequencies of the synchronized components.

It is important to notice that such a procedure lands itself nicely to study if and where cross-frequency synchronization is occurring in resting-state, when no specific task is taking place, and hence no hypothesis about the frequencies of CFC is available. Furthermore, this procedure only captures phase synchronization, since the amplitude does not affect the estimate [5]. This is of particular relevance, provided that a number of mechanisms are believed to operate simultaneously in the brain in order to allow communication between neuronal populations operating at different frequencies [4], but n:m synchronization is the only neuronal mechanism by which two neuronal population can influence each other at the temporal accuracy of the fast-operating neuronal population [22, 8]. Hence, the results provided by our procedure are interpreted in terms of a defined neurophysiological mechanism (i.e. n:m synchronization), and are very noise-resilient while being entirely independent from the amplitude of the signals. The detection of cross-frequency coupling is specifically relevant taking into account the phenomenon of frequency mixing, i.e. the appearance of new frequencies in neuronal circuitry when incoming oscillations are non-linearly integrated [51].

### 4.1 Rössler attractors

Firstly, we simulated synthetic data by using two Rössler attractors, as they retain non-linear properties that are similar to the ones displayed by real M/EEG data [52]. In order to simulate CFC, we modified one of the two attractors by applying a frequency shift. Similarly to what is shown in our previous work presenting the PLM [29], one can appreciate that the peak in the inter-ferometric spectrum grows monotonically as a function of the strength of synchronization between oscillators (regardless of their frequencies). Furthermore, one can appreciate that the interferometric spectrum peaks are at the frequency corresponding to the shift that had been introduced. Hence, the PSD allows the estimation of both the intensity of the synchronization as well as the difference between the frequencies of the involved signals. In addition, the resiliency to noise has been tested in this simulation, and a reliable estimate is possible also with realistic amount of noise.

### 4.2. Kuramoto oscillators

When dealing with real M/EEG signals, the case is more complicated since each signal has a very rich frequency spectrum, where the simultaneous presence of multiple components synchronizing in iso- and cross-frequency occurs. We used a model based on Kuramoto oscillators to explore if the CF-PLM can disentangle these different contributions.

Firstly, we explored the simple synchronization between two oscillators synchronized in iso-frequency (10 Hz-10 Hz). As shown by the peak centered in 0 in figure 5a, the synchronization is correctly captured as expected. Then, we explored the case of cross-frequency synchronization of Kuramoto oscillators. The interferometric spectrum displays one peak in correspondence to the frequency difference of the two oscillators (at −7 Hz in the example in 5b, since the two originating signals are oscillating at 10 Hz and 17 Hz, respectively). In the third case, one oscillator at 10 Hz has been compared to the sum of two more oscillators at 10 Hz and 17 Hz. This simulation is intended to create a single signal where some specific components are synchronized in iso-frequencies, while different ones are synchronized in cross-frequency. As expected, the components in iso-frequency produced a peak at 0 Hz in the interferometric spectrum, while a second peak appears at −7 Hz, capturing the cross-frequency synchronization. Such results show that the proposed methodology can disentangle the cases where multiple components are synchronized simultaneously in iso- and cross-frequency. Similarly to the previous scheme, we have then explored the resiliency of this estimate to the presence of noise. We show, also in this case, that our metrics produces a noise-resilient estimate of synchronization (despite being based solely on the phase).

Results reported in Figure 7 show that the proposed approach effectively estimates the frequencies involved in the coupling, both iso-frequency (Figure 7(a)), cross-frequency (Figure 7(b)) and simultaneous iso and cross-frequency (Figure 7(c)). More in details, the centers of all the computed crosses are correctly positioned and allow the identification of the frequency components present in the oscillators and involved in the coupling process.

### 4.3. Real Data

The analysis with the real data is intended to show that such a cross-frequency-coupling is happening in resting state, and can be captured by the proposed procedure. Each interferometric peak has been statistically validated against surrogates, making it unlikely that it would occur by chance. Furthermore, if the peaks were occurring by chance, one would expect that no consistent patterns of cross-frequency coupling in different subjects. However, observing the CFC analysis, one can appreciate the cross-frequency-coupling is not a widespread phenomenon happening in any area, but, rather, specific to some areas. More in detail, Figure 11, left column, clearly shows that the strongest cross-frequency connections do not spread randomly across the matrix, and a texture appears, indicating that a specific CFC network is operating involving specific edges, rather than being randomly spread across the brain, as one would expected for a random phenomenon. Besides this coherent spatial distribution, images in the right column of Figure 11 show that also the frequency components that are correlated are not random, but a pattern emerges. Finally, Figure 12 helps the visualization of the regions where cross frequency coupling occurs consistently. While a systematic description of these patterns goes beyond the scope of the current paper, one should notice that temporo-parietal regions as well as occipital ones, appear symmetrically involved in cross-frequency communication. Roughly speaking, these regions are involved in perceptive streams processing external stimuli. Importantly, as the ground truth is ultimately unknown, one should be very cautious at making inference.

**Figure 11:**
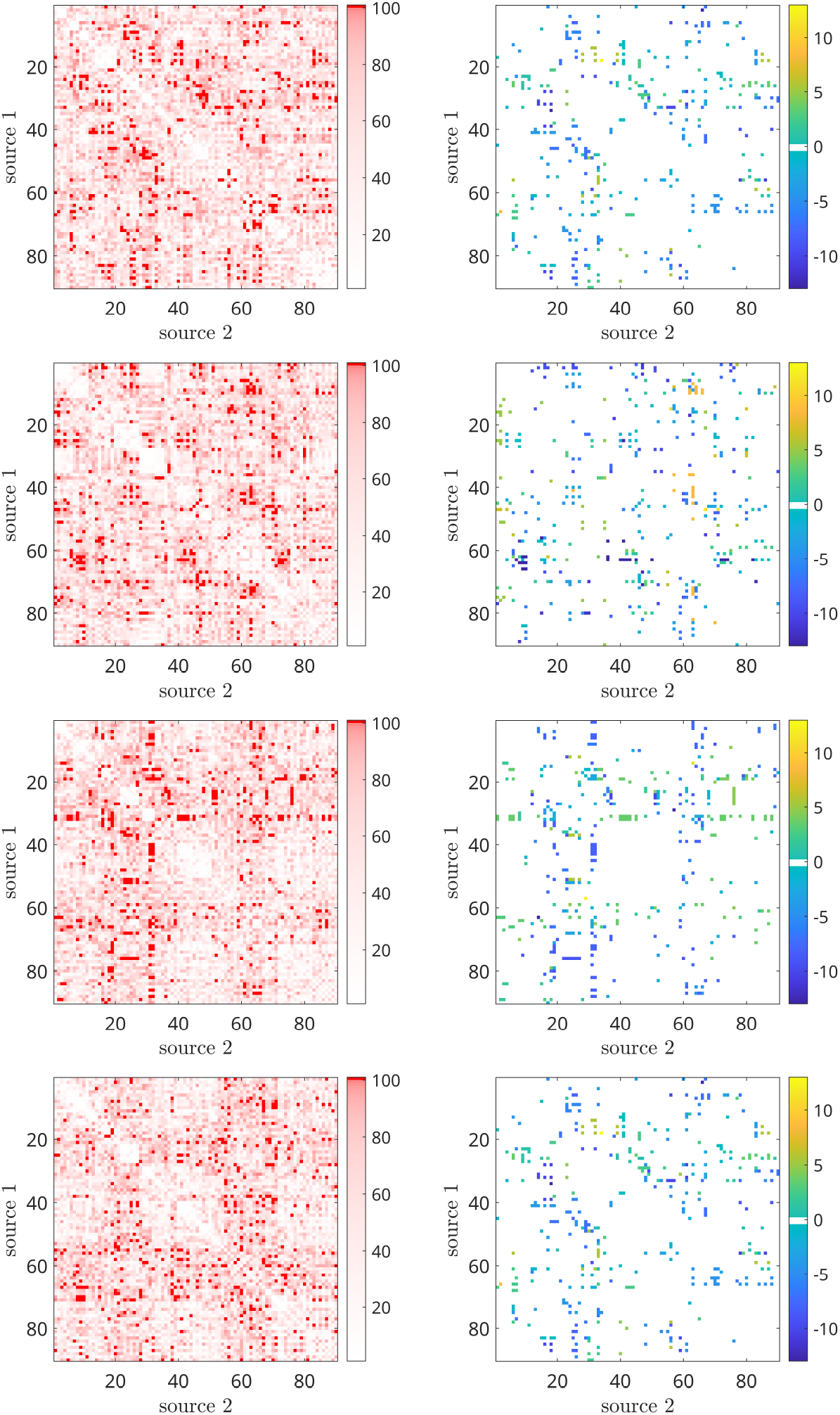
CFC matrix (left column: peak intensity, right column: peak frequency) of 4 subjects obtained by filtering the interferometric peaks intensity. The threshold values have been selected according to the percentiles of the distribution, spanning between 1 and 100. The most intense red points characterizes the CFC peaks with higher intensity.

**Figure 12:**
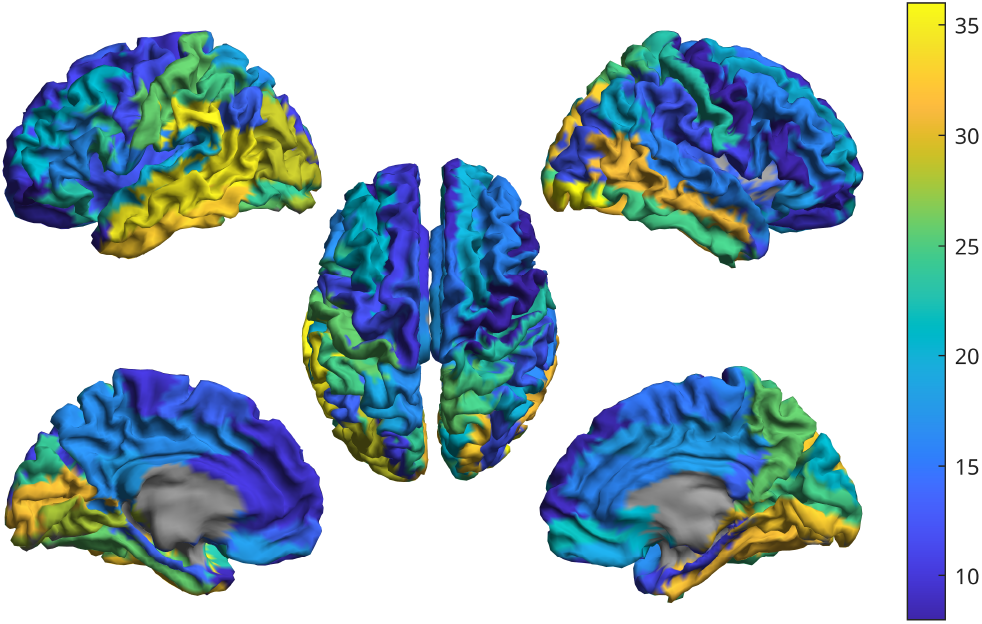
Brain regions where cross frequency coupling occurs consistently.

**Figure 13:**
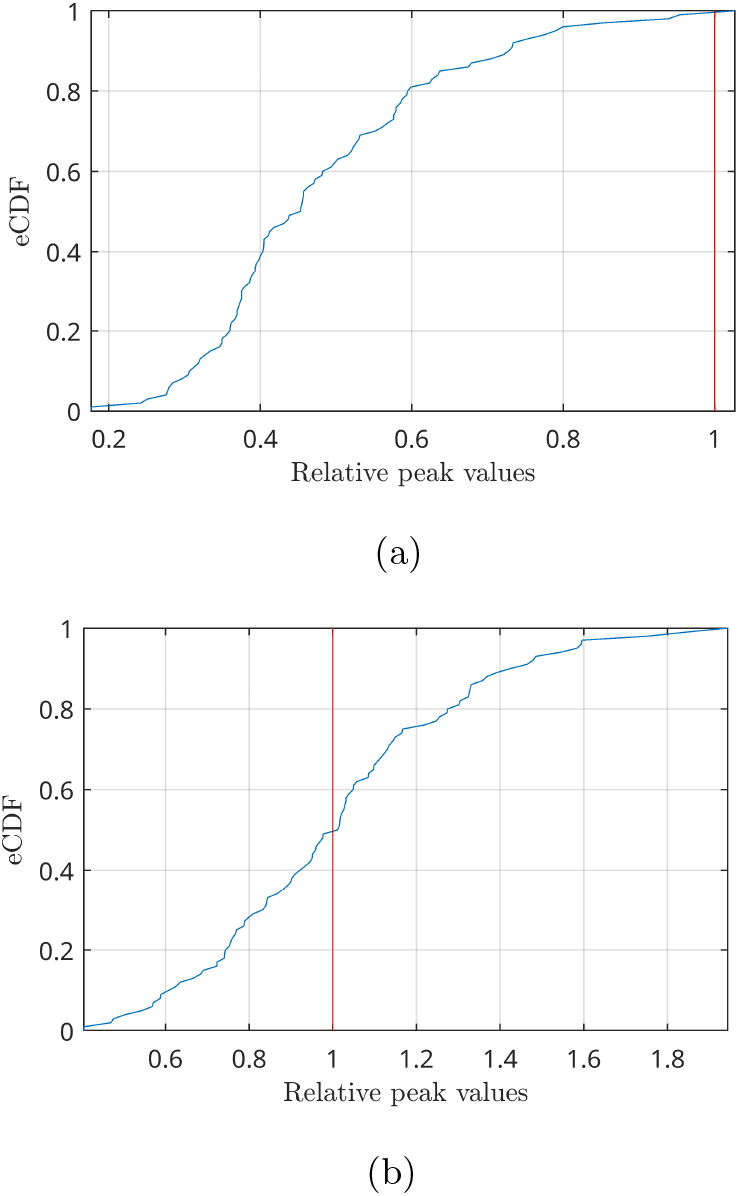
Empirical CDF of the peaks distributions obtained by shuffling the signal phases in the frequency domain (blue lines). Values have been normalized with respect to the highest CFC peak present in the data before shuffling (red line). The analysis refers to two cases: the highest CFC peak found among all couples of regions (a), where the peak intensity is above the 99% of the distribution, and a couple of regions with an average CFC peak intensity (b), where the peak is in the middle of the distribution.

However, one good aspect of the proposed procedure lies in the fact that a form of “double-check” is possible. In fact, once the frequency on the peak of the interferometric spectrum is known, and the related component in signal A has been identified, one has already a hypothesis about the frequency of the synchronized component in signal B. Hence, if the procedure is consistent, the filtering of signal B should confirm this hypothesis. In fact, as explained previously, the peak of the interferometric signal appears at a frequency equal to the difference of frequencies between the two components. In conclusion, with the proposed procedure we are able to determine the central frequencies of signals A and B involved in the coupling. We made a number of tests to explore the behaviour of the proposed procedure to different lengths of acquisitions and different number of epochs, confirming that this procedure is robust also with little and/or noisy data. One aspect of interest is that our metrics does not need the data to be split into trials, hence taking advantage of the full length of the available data. However, even when only short or limited data segments are available, the new procedure can still retrieve reliable results.

## 5. Conclusions

In this manuscript, we propose a new metric that can estimate cross-frequency coupling from broad-band signals with no a priori hypothesis on what the information transfers would be. Since cross-frequency coupling is the only neuronal mechanism that can allow fast communication between neuronal ensemble operating at different frequencies, we believe our metric can help to study the mechanisms of cross-frequency communication in the resting-state, as well as its topography and topology.

## Notes

### Competing Interest Statement

The authors have declared no competing interest.

